# The overall and sequence-specific degradation of soil extracellular DNA fragments: rates and influential factors

**DOI:** 10.64898/2026.01.14.699460

**Authors:** Ting Li, Song Zhang, Zelin Wang, Wei Huang, Zejin Zhang, Fang Wang, Dong Liu, Xiaoyong Cui, Rongxiao Che

## Abstract

While extracellular DNA persistence substantially influences soil microbiome investigations, its degradation kinetics remain poorly quantified. Here, we developed a primer-labeled DNA approach coupled with microcosm incubation to determine the overall and sequence-specific degradation rates of extracellular DNA amplicon fragments across China. We observed substantial variations in the overall degradation rates of extracellular 16S rRNA gene amplicon fragments among the study sites, with degradation rate constants ranging from 0.05 to 0.16 day^-1^. The overall degradation rate constants showed significant correlations with soil moisture content, prokaryotic abundance, prokaryotic community profiles, and mean annual precipitation (MAP). The significant influences of moisture content on the overall degradation rates were further verified by a moisture gradient microcosm experiment. The sequence-specific degradation rate constant profiles were additionally correlated with pH, nitrogen content, and mean annual temperature (MAT). Furthermore, propidium monoazide (PMA)-based exclusion of extracellular DNA signals significantly altered soil prokaryotic abundance, richness, and prokaryotic community profiles, and the pool sizes of sequence-specific extracellular 16S rRNA gene amplicon fragments were significantly correlated with their respective degradation rates. This study developed a methodology for determining the overall and sequence-specific degradation rates of extracellular DNA amplicon fragments, highlighting the profound influences of extracellular DNA on soil microbial research and informing the optimization of environmental DNA technologies.

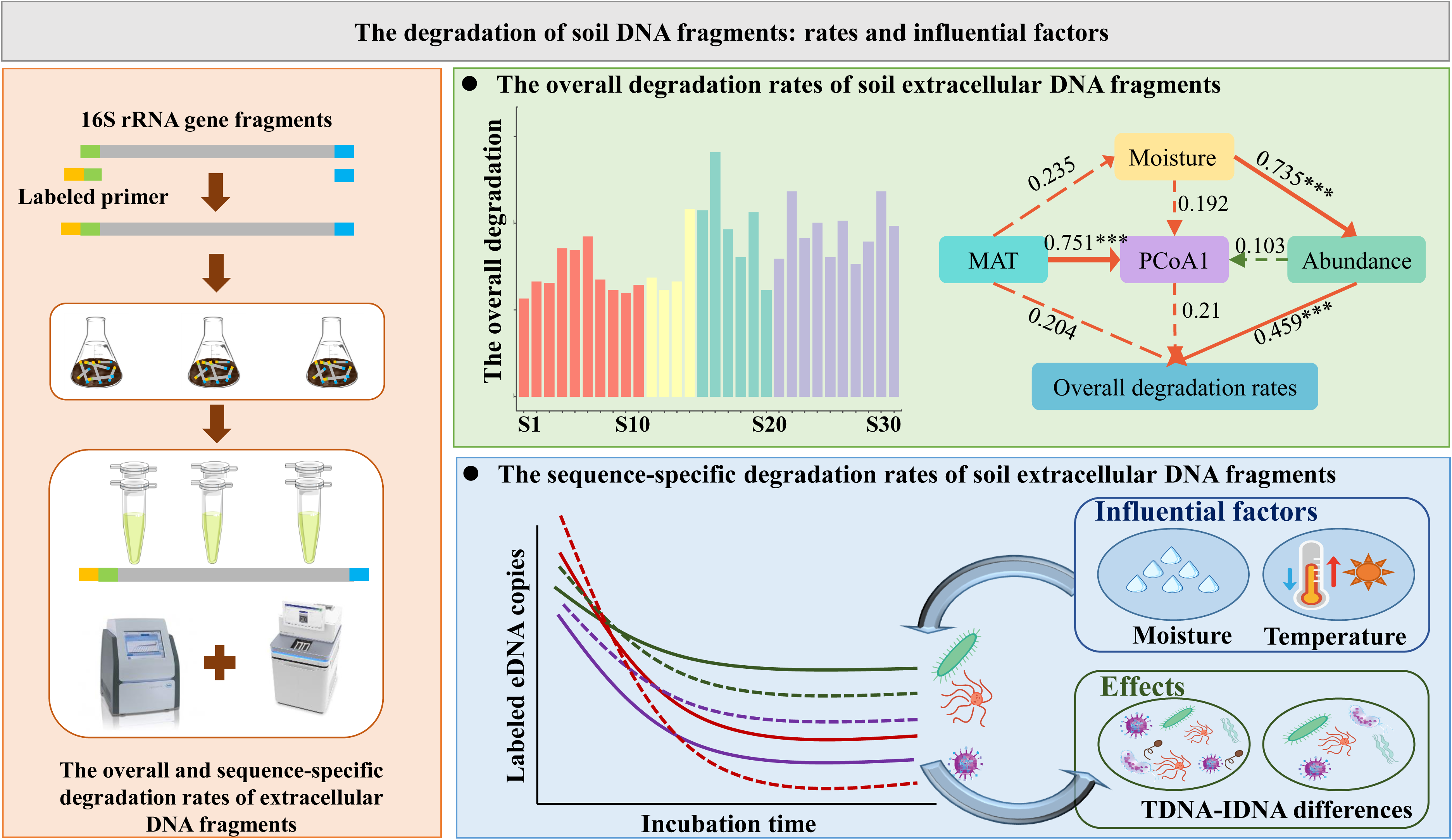

## 1. Introduction

The investigation of soil microbial abundance and diversity heavily relies on DNA-based technologies, such as real-time PCR, high-throughput amplicon sequencing, and metagenomic analysis (Che et al., 2018; Du et al., 2020; Yang et al., 2023). Soil DNA originates from both living and relic microbial cells, with the DNA from deceased microbial cells commonly referred to as relic or extracellular DNA (eDNA) (Ye et al., 2022). EDNA serves as a critical vector for horizontal gene transfer (HGT), facilitating the uptake of genetic material by competent microorganisms and promoting the spread of functional traits such as antibiotic resistance (Liu et al., 2024). In addition, eDNA participates in soil biogeochemical cycling because its enzymatic degradation releases bioavailable nutrients, particularly phosphorus and nitrogen, which can be reused by soil microorganisms (Ye et al., 2022). Moreover, the prevailing paradigm of total DNA extraction in soil microbiome studies introduces eDNA as a critical noise factor. Generally, its environmental persistence can lead to overestimation of microbial diversity in amplicon sequencing (Barnes et al., 2014; Wang et al., 2025; Xue et al., 2025) and distort qPCR-based quantification of marker genes (Carini et al., 2016; Sun and Ge, 2023; Wang et al., 2024a). A model simulation study showed that the extent of this influence is primarily determined by the abundance and sequence-specific degradation rates of eDNA (Lennon et al., 2018). Therefore, determining the overall and sequence-specific degradation rates of soil eDNA can provide crucial insights into evaluating the effects of eDNA on soil microbial analyses and the reliability of research based on environmental DNA technologies.

The overall degradation of soil eDNA have received significant attention due to their crucial roles in nutrient cycling, and horizontal gene transfer (Nagler et al., 2018). Early studies mainly assessed eDNA persistence and degradation using PCR, DNA hybridization, radioisotope labeling, and competent cell transformation techniques (Paget et al., 1992; Zhang et al., 2020; Samuels et al., 2025). These investigations consistently demonstrated that eDNA can persist in soils for months to years (Barnes et al., 2014; Pathan et al., 2020), maintaining its capacity to transform competent cells (Levy-Booth et al., 2007; Pietramellara et al., 2009). Most of these studies primarily concentrated on assessing the risks associated with transgenic technologies, narrowly examining the degradation dynamics of DNA sequences related to such technologies. Consequently, the insights they provided were usually qualitative or semiquantitative (Morrissey et al., 2015), limiting the comprehensive understanding of eDNA degradation dynamics. More recently, there has been a notable shift towards quantification, and the dynamics of soil eDNA were quantified in many studies (Eichmiller et al., 2016; Wei et al., 2018). For instance, the degradation rates of soil eDNA were quantified using real-time PCR with specific plasmid labels such as T7 and SP6 promoters (Ceccherini et al., 2009). Another study simultaneously determined the dynamics of both soil microbial community and eDNA by adding 16S rRNA gene primer labels to exogenous DNA (Sirois and Buckley, 2019). Moreover, stable isotopic probing was also utilized for determining the degradation rates of soil eDNA (Morrissey et al., 2015). However, most quantitative approaches have remained restricted to single or highly specific DNA targets (*e.g*., transgenic sequences), thereby overlooking sequence-specific variation in soil eDNA degradation rates (Pietramellara et al., 2009; Sirois and Buckley, 2019; Wang et al., 2019).

In this study, “sequenceLspecific degradation” refers to statistically significant differences in firstLorder degradation rate constants (k, dayL¹) among distinct 16S rRNA gene amplicon sequence variants (ASVs) under identical soil and incubation conditions. The potential variations in sequence-specific eDNA degradation rates can be attributed to several factors. First, sequence-dependent degradation can arise from differences in nucleotide composition, particularly GC content. This influences the thermodynamic stability and base-stacking interactions of the DNA duplex, thereby altering its accessibility to extracellular nucleases (Marrone and Ballantyne, 2008; Wolpe and Guertin, 2022). Second, local conformational features and the formation of potential secondary structures, such as stem-loops or hairpins, can create steric hindrance that protects the phosphodiester backbone. Differences in base composition also alter the elemental stoichiometry (*e.g*., C:N ratio) of DNA molecules, potentially affecting microbial preference for recycling specific sequences as nutrient sources (Cai et al., 2006a; Buitrago et al., 2021). Third, the persistence of soil DNA can be mainly attributed to its adsorption and protection by minerals and humus in soils (Cai et al., 2006b; Vuillemin et al., 2017; McKinney and Dungan, 2020). Thus, sequence-dependent differences in the physicochemical behavior of DNA molecules, including their affinity for soil minerals and organic matter, may also contribute to variation in degradation rates among sequences (Levy-Booth et al., 2007; Morrissey et al., 2015). Consequently, we proposed three central hypotheses. (1) The degradation rates of eDNA amplicon fragments were expected to be highly sequenceLspecific. (2) The rates and patterns of eDNA fragments degradation would be influenced by environmental factors such as temperature and moisture content. (3) The sequenceLspecific degradation of extracellular 16S rRNA gene amplicon fragments would significantly influence estimates of soil prokaryotic abundance and diversity.

To test these hypotheses, we investigated the overall and sequence-specific degradation rates of soil extracellular 16S rRNA gene amplicon fragments and the factors influencing these rates. Additionally, the effects of extracellular 16S rRNA genes on soil prokaryotic community analysis and their relationships with the overall and sequence-specific degradation rates were determined. Soil samples were collected from 30 representative ecosystems across China. A new methodology, involving the addition of exogenous 16S rRNA gene amplicon fragments tagged with specific primers, microcosm incubation, real-time PCR, and amplicon sequencing, was developed to determine the overall and sequence-specific degradation rates of soil extracellular 16S rRNA gene amplicon fragments. We employed 16S rRNA gene amplicon fragments as the standardized and trackable model substrate because the 16S rRNA gene is one of the most extensively used molecular markers in microbiome research (Knight et al., 2018; Du et al., 2025).

## 2. Results

### 2.1. The overall degradation rate constants of soil extracellular 16S rRNA gene amplicon fragments

The glyceraldehyde-3-phosphate dehydrogenase (GAPDH F)Ltagged 16S rRNA gene amplicon fragments were consistently detectable throughout the 48-day incubation period, but their abundance rapidly declined as the incubation progressed (Fig. 1a, *P* < 0.05). After 48 days of incubation, 0.2–3.1% of the initially spiked GAPDH FLtagged 16S rRNA gene amplicon fragments persisted in the soils (Fig. 1a). The degradation rate constants of the spiked extracellular 16S rRNA gene amplicon fragments displayed considerable variability among the study sites, ranging from 0.05 to 0.16 day^-1^ (Fig. 1b). Furthermore, we found that degradation rate constants differed significantly among ecosystem types (Fig. 1c, *P* < 0.05). Specifically, cropland and forest soils exhibited significantly higher degradation rates than grassland soils (*P* < 0.05). Random forest modeling revealed that soil moisture content, prokaryotic abundance, and prokaryotic community profiles were critical predictors for the overall degradation rate constants of extracellular 16S rRNA gene amplicon fragments, and they all showed significant positive correlations (Figs. 1d and S1a–d). However, we did not observe significant correlations between soil texture and the degradation rate constants (Fig. S1e and f). Structural equation modeling (SEM) analysis indicated that the overall degradation rate constants of 16S rRNA gene amplicon fragments were mainly directly affected by prokaryotic abundance, which was indirectly influenced by soil moisture (Fig. 1e). In the moisture gradient microcosm experiment, we also observed a strong positive correlation between soil moisture content and the overall degradation rates of extracellular 16S rRNA gene amplicon fragments (Fig. 1f).

**Fig. 1.**
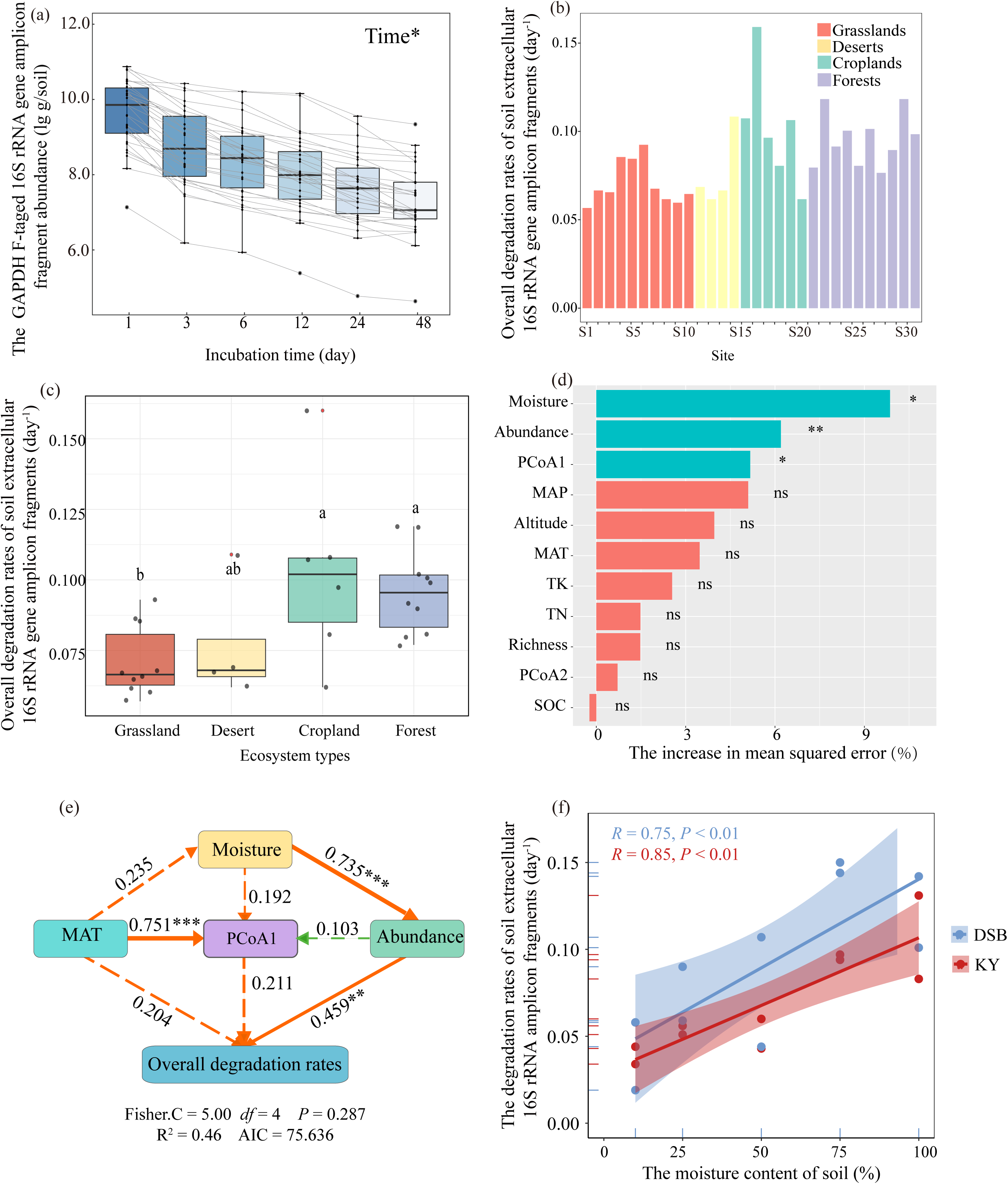
The overall degradation rates of soil extracellular 16S rRNA gene amplicon fragments and their influential factors. (a) The GAPDH F-tagged 16S rRNA gene amplicon fragment abundance at different incubation time points. (b) and (c) The degradation rate constants of soil extracellular 16S rRNA gene amplicon fragments across the study sites and different ecosystem types. (d) The factors influencing extracellular 16S rRNA gene degradation rates. (e) The influencing factors for the overall degradation rates of soil extracellular 16S rRNA gene amplicon fragments revealed by structural equation modeling. Orange and green lines indicate positive and negative relationships, respectively. Solid and dashed lines indicate significant and non-significant relationships, respectively. Path coefficients are denoted by numbers adjacent to the arrows, with arrow width reflecting their strength. (f) The influence of soil moisture on the degradation rates of soil microbial eDNA. Significance levels are indicated as follows: * *P* < 0.05, ** *P* < 0.01, and *** *P* < 0.001. Moisture: soil moisture content; Abundance: prokaryotic abundance; NMDS1: the scores at the first axis of the NMDS ordination of prokaryotic community profiles; MAP: mean annual precipitation; Richness: soil prokaryotic richness; NMDS2: the scores at the second axis of the NMDS ordination of prokaryotic community profile; TK: soil total potassium contents; MAT: mean annual temperature; AP: soil available phosphorus content; TN: soil total nitrogen contents; and TOC: soil total organic carbon content.

### 2.2. The sequence-specific degradation rate constants of the extracellular 16S rRNA gene amplicon fragments

The richness of the GAPDH FLtagged 16S rRNA gene amplicon fragments significantly decreased during the incubation period, declining to approximately 60% of the initial values after 48 days (Fig. 2a). The prokaryotic community profiles based on the GAPDH FLtagged 16S rRNA gene amplicon fragments also exhibited significant variations across different incubation time points (Figs. 2b and S2), and community-profile similarities declined more strongly with increasing incubation intervals (Fig. S3). These findings suggest sequence-specific differences in degradation patterns among ASVs. Indeed, the degradation rate constants for different extracellular 16S rRNA gene amplicon fragments were within 0.40 day^-1^, with most ranging between 0.06 and 0.12 day^-1^ (Fig. 2c). For example, many sequences assigned to Proteobacteria had significantly higher degradation rate constants than those assigned to Acidobacteriota, whereas Methylomirabilota showed intermediate degradation rate constants (Fig. 3a). The sequence-specific degradation rate constant profiles of the extracellular 16S rRNA gene amplicon fragments also showed significant correlations with multiple environmental factors, including soil moisture, mean annual temperature (MAT), mean annual precipitation (MAP), soil pH, as well as the contents of soil ammonium nitrogen (NH4^+^-N) and nitrate nitrogen (NO_3_L-N) (Fig. 3b). However, no significant relationships were found between the degrdation rates and the GC content of the representative sequences (Fig. S4). Further analysis suggested that the degradation rate constants of extracellular 16S rRNA gene amplicon fragments of the dominant phyla showed similar correlations with the environmental factors, and MAT was identified as the strongest predictor for them (Fig. S5).

**Fig. 2.**
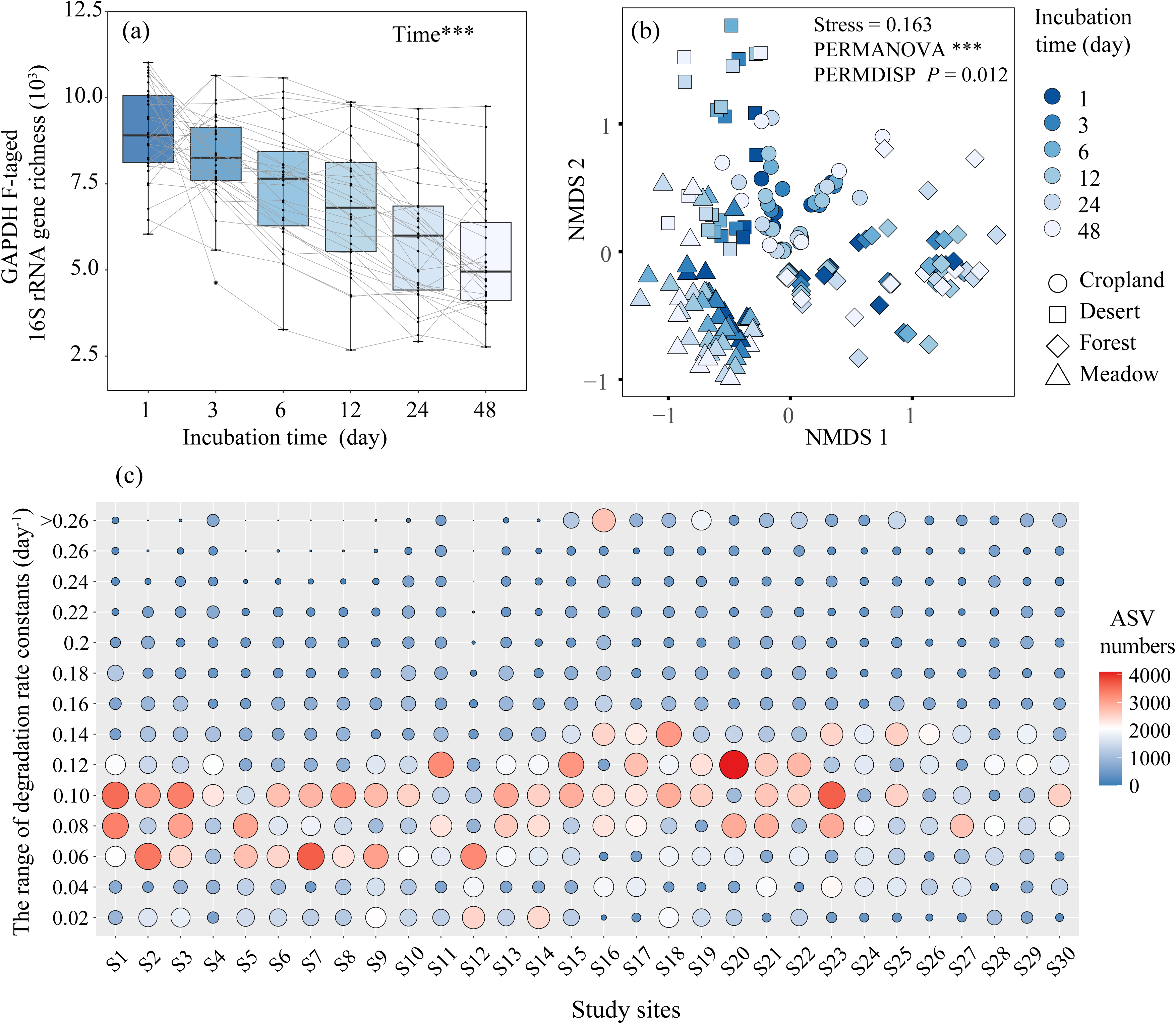
The sequence-specific degradation rates of soil exogenous extracellular 16S rRNA gene amplicon fragments. (a) Soil GAPDH F-tagged 16S rRNA gene richness. (b) The nonmetric multidimensional scaling (NMDS) ordination of the community profiles based on GAPDH FLtagged 16S rRNA gene amplicon fragments at different incubation time points. (c) The number of GAPDH F-tagged 16S rRNA gene ASVs within different degradation rate ranges.

**Fig. 3.**
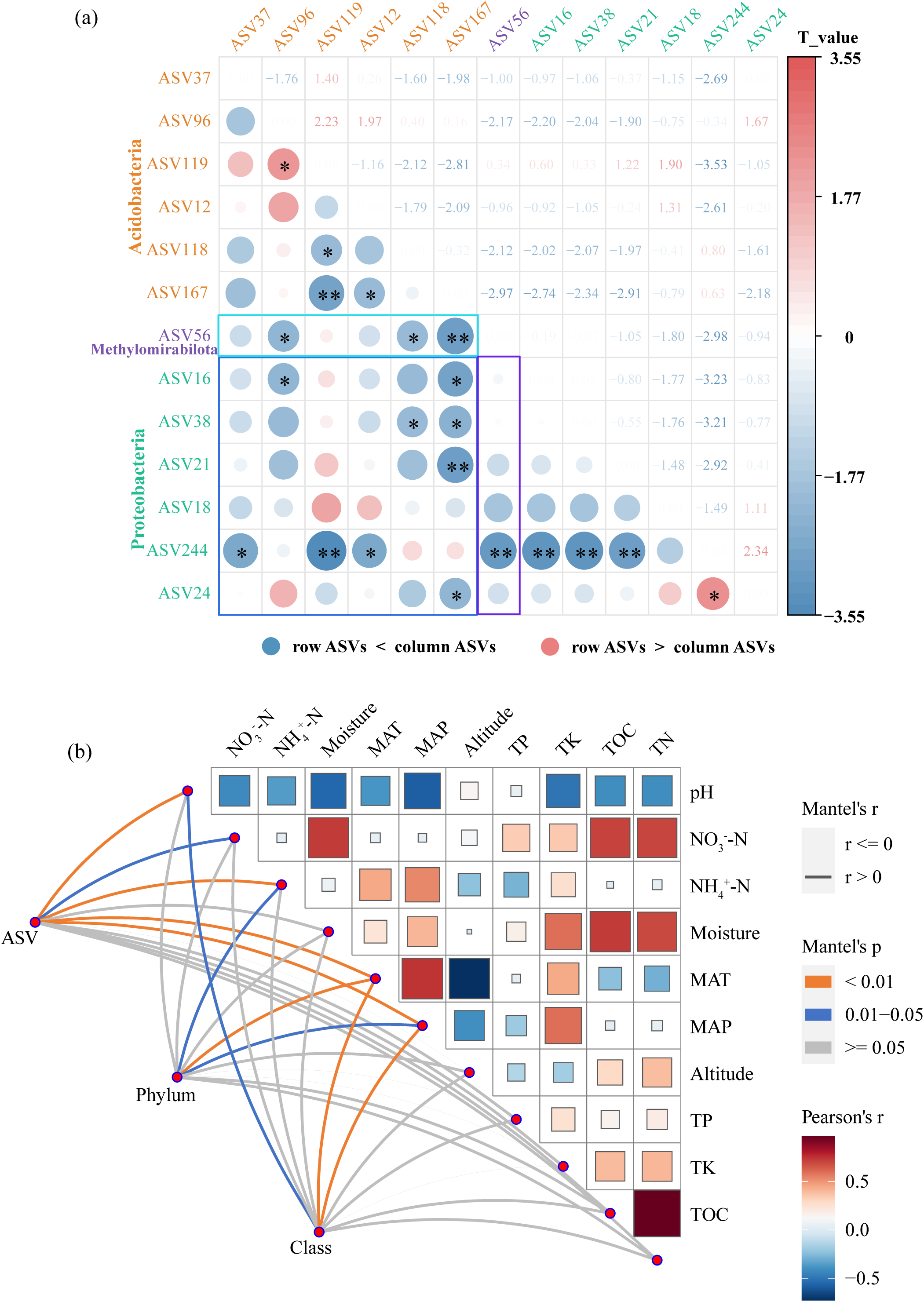
The difference in sequence-specific degradation rates of soil extracellular 16S rRNA gene amplicon fragments and their influencing factors. (a) The paired comparison of degradation rates among different ASVs. In the heatmap, each cell represents a pairwise comparison between two ASVs. Blue indicates that the degradation rate of the ASVs listed in the row (row ASVs) is significantly lower than that of the ASVs listed in the column (column ASVs); red indicates that the row ASVs has a significantly higher degradation rate than the column ASV. A positive *t* value indicates that the row ASVs degrades significantly faster than the column ASVs; a negative *t* value indicates the opposite. Significance levels are indicated as follows: * *P* < 0.05, ** *P* < 0.01, and *** *P* < 0.001. (b) The relationships between the sequence-specific degradation rate profiles and environmental factors. NO_3_^-^-N: soil NO_3_^-^-N contents; NH_4_^+^-N: soil NH_4_^+^-N contents; TN: soil total N contents; TP: soil total P contents; TK: soil total K contents; AP: soil available P contents; TOC: soil total organic carbon content; MAT: mean annual temperature; and MAP: mean annual precipitation.

### 2.3. The influences of extracellular 16S rRNA genes on soil prokaryotic community analysis and their links with the degradation rates

Based on the PMA treatment, we observed significant effects of extracellular 16S rRNA genes on the analysis of soil prokaryotic abundance and diversity (Figs. 4 and 5). The PMA treatment revealed that intact cells accounted for approximately 40% (range: 9–73%) of the total 16S rRNA gene copies. In contrast, over 80% (range: 27–97%) of the observed ASV richness was associated with sequences originating from intact cells (Fig. 4a and b). Meanwhile, the Shannon index of the PMA-treated prokaryotic communities was significantly lower than that of the total prokaryotic community (Fig. 4c). Furthermore, significant differences were found between the profiles of total and PMA-treated prokaryotic communities, especially in the forest and cropland ecosystems (Figs. 4d–f and S6). The Bray-Curtis dissimilarity between the intact cells and the total prokaryotic community was approximately 52.8% (Fig. 4f). For instance, Abditibacteriota, Bacteroidota, Nitrospirota, Fibrobacterota, Entotheonellaeota, Elusimicrobiota, and Armatimonadota were significantly enriched in the total prokaryotic community, whereas Actinobacteriota and Planctomycetota showed opposite trends (Fig. 5a). However, many other microbial taxa, such as Proteobacteria, Acidobacteriota, and Chloroflexi showed no significant differences between the total and PMA-treated prokaryotic communities (Fig. 5a). Additionally, the correlations with environmental factors were stronger for the total soil prokaryotic community structure than the PMA-treated community structure (Fig. 5b). Total prokaryotic community structure was significantly correlated with soil pH, total potassium (TK), NH_4_^+^-N, MAT, and MAP. However, the PMA-treated prokaryotic community structure was only significantly correlated with soil pH, MAT, and MAP (Fig. 5b).

**Fig. 4.**
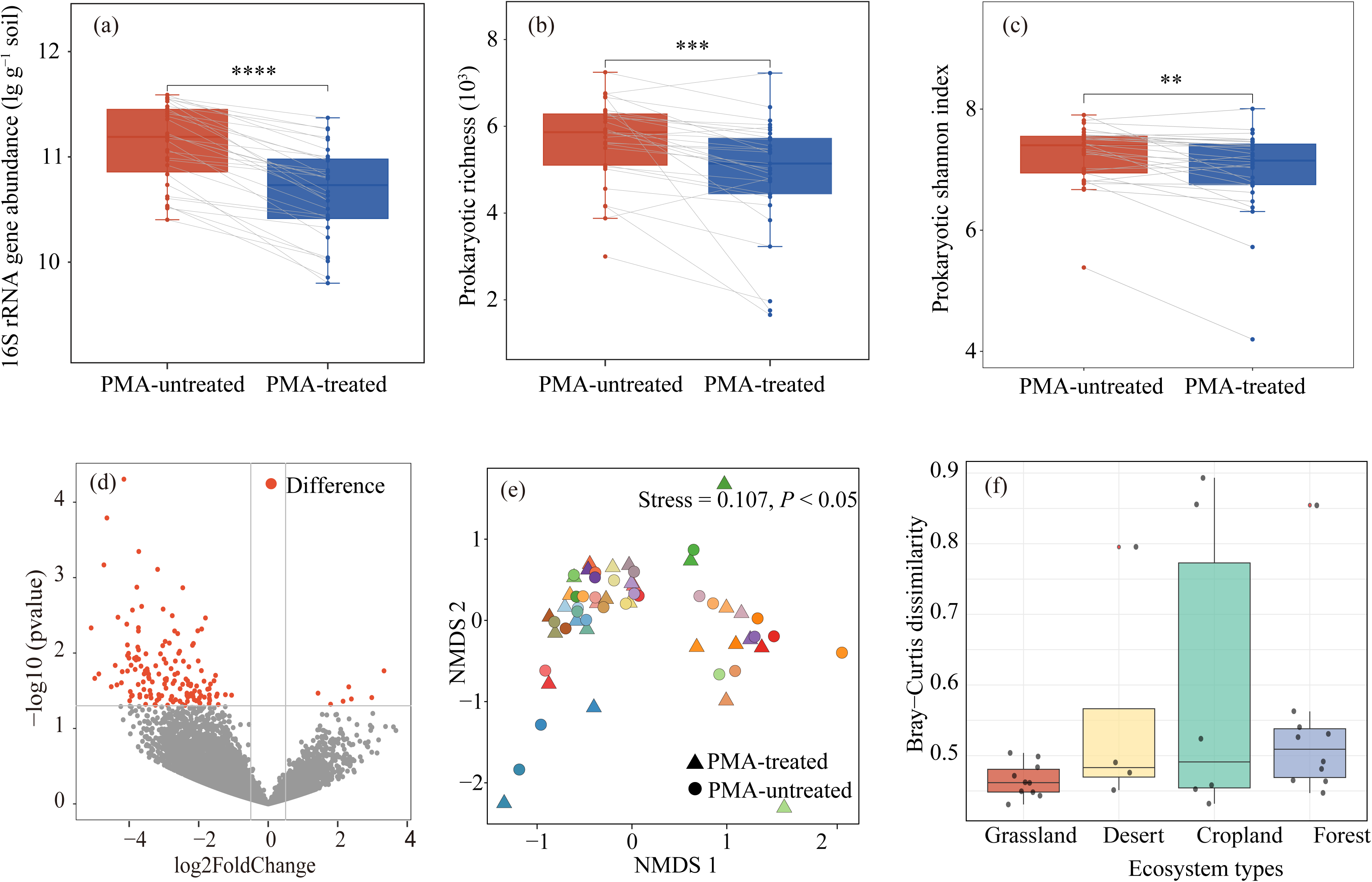
The differences in the abundance, richness, Shannon, and community composition between total and PMA-treated soil prokaryotes. (a) The abundance of total and PMA-treated soil prokaryotes. (b) and (c) The richness and Shannon index of total and PMA-treated soil prokaryotes. (d) The differences between the relative abundance of total and intracellular ASVs. The red points represent the prokaryotic taxa exhibiting statistically significant differences. (e) The nonmetric multidimensional scaling (NMDS) ordination of total and PMA-treated soil prokaryotes. Different colors represent samples from different sites. (f) The Bray-Curtis dissimilarity between total and PMA-treated soil prokaryotes.

**Fig. 5.**
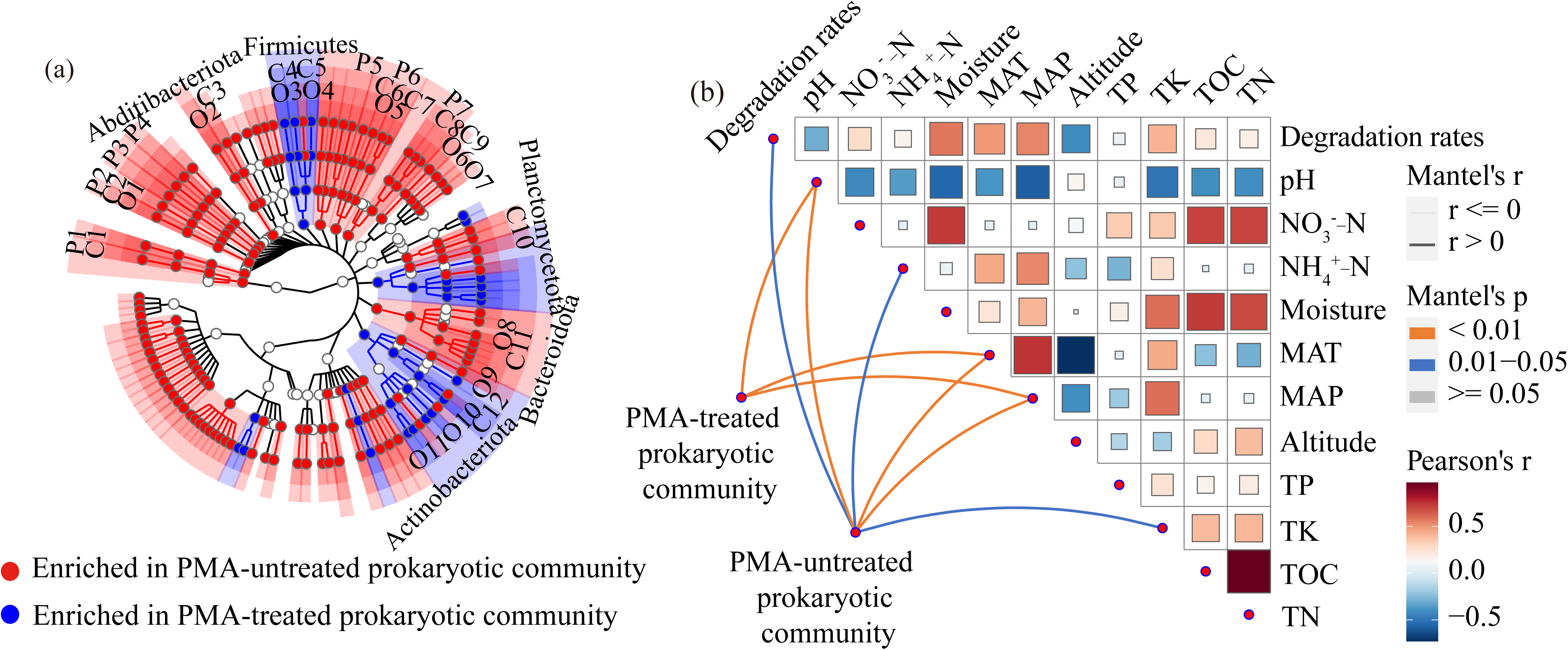
Effects of eDNA exclusion on soil prokaryotic community composition and its relationships with environmental factors. (a) The taxa with significant differences between total and intact cell prokaryotic communities. P1: Thermoplasmatota; P2: Nitrospirota; P3: Fibrobacterota; P4: Entotheonellaeota; P5: Elusimicrobiota; P6: Armatimonadota; P7: Myxococcota; C1: Thermoplasmata; C2: Nitrospiria; C3: Vampirivibrionia; C4: Clostridia; C5: Bacilli; C6: Armatimonadia; C7: Chthonomonadetes; C8: Polyangia; C9: Blastocatellia; C10: Parcubacteria; C11: Bacteroidia; C12: Thermoleophilia; O1: Nitrospirales; O2: Abditibacteriales; O3: Clostridiales; O4: Bacillales; O5: Armatimonadales; O6: Pyrinomonadales; O7: Bryobacterales; O8: Chitinophagales; O9: Rubrobacterales; O10: Propionibacteriales; and O11: Corynebacteriales. (b) The relationships between soil prokaryotic community profiles and environmental factors. NO_3_^-^-N: soil NO_3_^-^-N contents; NH_4_^+^-N: soil NH_4_^+^-N contents; Moisture: soil moisture content; MAT: mean annual temperature; MAP: mean annual precipitation; TP: soil total P contents; TK: soil total K contents; and TOC: soil total organic carbon contents.

Interestingly, a significant negative correlation was observed between the total prokaryotic community structure and the degradation rates of extracellular 16S rRNA gene amplicon fragments. However, no significant relationship was observed for the PMA-treated prokaryotic communities (Fig. 5b). The relationships between extracellular 16S rRNA gene pool sizes and degradation rates were further explored. We found a significant positive correlation between the overall degradation rates and the copies of soil extracellular 16S rRNA genes (Fig. S1c). Moreover, most study sites exhibited significant positive correlations between sequence-specific degradation rates and the pool sizes of soil extracellular 16S rRNA genes (Fig. S1g). Additionally, there were significant correlations between the differences in relative abundance of taxa in the total and PMA-treated prokaryotic community and the sequence-specific degradation rates; but these relationships varied across the study sites (Fig. S1g).

## 3. Discussion

In this study, we found that extracellular 16S rRNA gene amplicon fragments persisted in soils for at least several weeks and the degradation rate constants ranged from 0.05 to 0.16 day^-1^. The degradation rate constants of soil eDNA based on plasmid and stable isotope labelling microbial genomes were usually 0.03–0.14 day^-1^ (Morrissey et al., 2015; Sirois and Buckley, 2019; Wang et al., 2019), which is consistent with our results. Therefore, the PCRLgenerated DNA fragments provide a standardized substrate for quantifying the degradation kinetics of added DNA under controlled conditions, offering a useful proxy for eDNA turnover. Additionally, substantial variations in the overall degradation rates of added exogenous eDNA amplicon fragments were also observed in this study, which can be mainly elucidated in the following ways (Fig. 1a and b). First, the degradation of soil eDNA is strongly influenced by the availability of enzymes (Nihemaiti et al., 2020). As a large proportion of soil enzymes originate from microbes (Bhardwaj et al., 2024; Tan et al., 2025), differences in microbial abundance among study sites could make a substantial contribution to the variations in the overall degradation rates. Second, different soil microbial taxa play diverse roles in eDNA degradation. In this study, there were considerable discrepancies in soil community profiles across study sites, providing another plausible explanation for the differing degradation rates of eDNA. Third, environmental factors, including soil moisture, pH, and temperature, can predominently govern enzymatic reaction rates (He et al., 2024; Shah et al., 2024). Indeed, strong positive correlations were observed between moisture content and eDNA degradation rates in both the survey and microcosm experiments (Fig. 1d–f). These results support our second hypothesis that environmental factors regulate the degradation rates and persistence patterns of eDNA fragments, with soil moisture emerging as a particularly important driver. This finding further suggests that dryland soils may be particularly susceptible to eDNA persistence, potentially increasing the risk of overestimating microbial abundance and diversity in arid ecosystems (Carini et al., 2016; Lennon et al., 2018).

Consistent with our first hypothesis, we observed that the degradation of soil eDNA amplicon fragments showed strong sequence-specific patterns (Fig. 2c). In particular, many sequences belonging to Proteobacteria exhibited a significant faster DNA degradation rate than those of Acidobacteriota (Fig. 3a). The sequence-specific degradation of the eDNA amplicon fragments can be interpreted from multiple perspectives. First, the variation in the number and position of restriction enzyme cutting sites for different sequences could be a critical reason for the sequence-specific degradation rates of eDNA (Brown, 2020). Second, sequence differences could also influence the spatial structure and mineral adsorption of DNA fragments (Cleaves II et al., 2011; Buitrago et al., 2021), indirectly affecting the sequence-specific degradation rates. Third, microbes might preferentially degrade and recycle the abundant DNA sequences present in their habitats. Thus, the sequence-specific degradation rates could also be ascribed to the varied abundance of different eDNA amplicon fragment sequences (Finkel and Kolter, 2001). This assertion is further evidenced by the generally observed positive correlations between eDNA abundance and degradation rates (Fig. S1). We also examined whether GC content could explain the observed sequenceLspecific patterns, but no significant correlation was found (Fig. S4), suggesting that simple base composition is not the primary driver in this study. However, this does not exclude the possibility that higherLorder structural features (*e.g*., hairpin loops) or sequenceLspecific nuclease recognition motifs contribute to differential degradation (Wang et al., 2007). This should be tested in future studies using synthetic DNA constructs with controlled structural elements. As observed in this study, the differential sequence-specific degradation rates also led to significant alterations in the community profiles based on the exogenous extracellular 16S rRNA genes (Fig. 2a and b). Additionally, this study essentially determined the effects of DNA sequences on the extracellular 16S rRNA gene amplicon fragments degradation rates. However, actual sequence-specific degradation rates of extracellular 16S rRNA genes can be additionally influenced by the varying degrees of protection offered by cell residuals from different microbial taxa (Shi et al., 2024). Thus, the differences in the actual sequence-specific degradation rates of extracellular 16S rRNA genes should be even more significant than those observed in this study. This can be one of the main reasons for the influences of eDNA on the analysis of soil microbial community profiles (Lennon et al., 2018).

Accordingly, we further explored how eDNA may influence prokaryotic community analyses using PMA treatment, and significant disparities were observed between the profiles of the total and PMA-treated soil prokaryotic communities (Fig. 4). The differences between total and PMA-treated community profiles can arise from multiple factors. As mentioned above, the disparities can be ascribed to the differential sequence-specific degradation rates of eDNA. Additionally, these differences are also related to several other factors. The primary reason should be the distinct source of extracellular and intracellular DNA which are derived from dead and live microbes, respectively. The death of microorganisms is mainly caused by environmental selection or stochastic processes (Zhang et al., 2016). In terms of environmental selection, the dead microbial taxa should be less competitive than PMA-treated ones (Upton et al., 2019; Chu et al., 2020). The stochastic death should be correlated with the population size of each microbial species, but the varied stochastic mortality among different microbial taxa can still lead to different community profiles between dead and live microbes (Blazewicz et al., 2020). This study also revealed that extracellular 16S rRNA genes led to substantial overestimation of soil prokaryotic abundance and diversity (Figs. 4 and 5). These findings were supported by many recent studies (Carini et al., 2016; Sun and Ge, 2023; Du et al., 2025) and can be explained as follows. The overestimated prokaryotic abundance can be attributed to the long-term persistence of eDNA (Sun and Ge, 2023). However, as DNA extraction efficiency may differ between intact cells and eDNA, the actual differences between total and living prokaryotic abundance could be smaller than those observed in this study. Similarly, the overestimated prokaryotic richness may arise from historically accumulated microbial taxonomic information stored in eDNA pools (Deshpande and Fahrenfeld, 2023; Wang et al., 2024b).

We observed a significant correlation between eDNA degradation rates and the overall structure of the prokaryotic community, but this relationship was absent in PMA-treated communities (Fig. 5b). This discrepancy highlights the divergent ecological roles of extracellular and intracellular DNA. Analyses of the total community integrate intracellular DNA from metabolically active cells with eDNA which primarily originates from historical microbial residues (Lennon et al., 2018). EDNA incorporates signals that likely reflect the legacy effects of past environmental conditions (Wang et al., 2021). In contrast, the PMA-treated community reflects transient microbial activity driven by current selective pressures. Additionally, eDNA can serve as a nutrient source and facilitate horizontal gene transfer, which may further shape its interactions with contemporary microbial communities (Levy-Booth et al., 2007).

Notably, we also observed significant positive correlations between sequence-specific degradation rates and the pool sizes of extracellular 16S rRNA gene amplicon fragments at most of the study sites (Fig. S1g). This finding suggests that abundant eDNA degrades at a faster rate compared to rare eDNA. As mentioned earlier, this could be explained by several mechanisms. First, as soil eDNA is subject to enzymatic degradation and microbial recycling, abundant DNA sequences may be more likely to be encountered and degraded by extracellular nucleases simply due to their higher copy numbers (Levy-Booth et al., 2007; Nagler et al., 2018). Similarly, if microbes preferentially take up DNA as a nutrient source, they may degrade abundant sequences more frequently as a stochastic consequence of higher encounter rates (Finkel and Kolter, 2001). However, we also found that the relationships between the sequence-specific degradation rates and the effect sizes of extracellular 16S rRNA gene amplicon fragments varied across the study sites (Fig. S1g). The sequence-specific effect sizes of extracellular 16S rRNA gene amplicon fragments are mainly determined by both their production and degradation rates (Pietramellara et al., 2009; Sirois and Buckley, 2019). These inconsistent correlations emphasize the critical role played by the production rates of extracellular 16S rRNA genes in influencing the analysis of prokaryotic communities. Therefore, future studies should systematically\ determine both the production and degradation rates of eDNA.

Despite the high-resolution insights afforded by our methodology, several limitations should be considered. First, utilizing PCR-amplified 16S rRNA gene fragments as proxies oversimplifies the structural and sequence complexity of natural soil eDNA pools. In natural environments, eDNA varies widely in fragment length and conformation, and exhibits complex interactions with mineral surfaces, all of which fundamentally affect degradation dynamics (Levy-Booth et al., 2007; McKinney and Dungan, 2020). Additionally, the highly conserved nature of the 16S rRNA gene means that the nucleotide variability explored here (*e.g.*, GC content gradients) does not fully capture the genomic heterogeneity of entire metagenomes (Knight et al., 2018). Consequently, our reported degradation rates indicate the decay potential of highly accessible linear eDNA rather than a universal rate for all soil DNA fractions. Future studies incorporating diverse metagenomic DNA, especially those with extreme AT or GC contents, are essential for building a more generalizable predictive framework for eDNA persistence (Morrissey et al., 2015). Second, methodological biases inherent in quantifying the intracellular community must be acknowledged (Du et al., 2025). Although PMA treatment is widely used to exclude eDNA, its efficiency in complex soil matrices can be compromised by limited light penetration in turbid suspensions and competitive adsorption to soil particles (Nocker et al., 2007; Carini et al., 2016; Heise et al., 2016). Compounding this issue, downstream DNA recovery is subject to differential cell lysis, as taxa with robust cell walls (*e.g.*, Gram-positive bacteria) may resist extraction (Frostegård et al., 1999; Albertsen et al., 2015). While our standardized bead-beating protocol and calculation of degradation rate constants (*k*) minimize systematic biases, future studies should integrate complementary viability markers (*e.g.*, RNA-based analyses or protein synthesis activity probes) and multi-extraction comparisons to robustly validate these ecological patterns (Emerson et al., 2017).

## 4. Materials and methods

### 4.1. Study sites and soil sampling

Soil samples were collected from 30 sites across China (Fig. S7). The selection of sampling sites was mainly based on National Soil Fertility and Fertilizer Effect Long-term Monitoring Network and the Chinese Ecosystem Research Network (CERN). Several other sampling sites were included, based on the systematic consideration of ecosystem typicality, soil types, and climates. The 30 sites spanned major climatic zones and land-use types in China, including 10 grasslands, 10 forests, 6 croplands, and 4 deserts, ensuring broad applicability of the results across diverse environmental scenarios. The longitude of the sites ranged from 80.724 °E to 124.817 °E, while the latitude ranged from 21.917 °N to 50.168 °N. The altitude of the study sites varied from 13 m to 4397 m. The ranges of mean annual temperature (MAT) and precipitation (MAP) were −3.35–22.53L and 39–1809 mm, respectively. The geographic and climate information of study sites is detailed in Table S1.

At each sampling site, 10 subsampling points were randomly selected, with a minimum distance of 10 m between adjacent subsampling points. Soil samples (0–20 cm) from 10 subsampling points at each study site were collected, thoroughly homogenized, and sieved to ≤ 2 mm to form a composite sample. Subsequently, all the composite soil samples were divided into two sub-samples. The first sub-sample was air-dried for the determination of soil pH values, as well as measurements of total organic carbon (TOC), total nitrogen (TN), total phosphorus (TP), and available phosphorus (AP). The second sub-sample was preserved at 4°C for the determination of moisture content, inorganic nitrogen content, as well as for microcosm experiment and PMA treatment.

### 4.2. Analysis of soil physicochemical properties

Soil moisture content was determined by drying the soils at 105 °C for 48 hours (Che et al., 2019). Soil pH values were measured using a pH meter (Mettler Toledo, Switzerland) with a soil-to-water ratio of 1:2.5 (Zhang et al., 2023a). TOC content of the soils was examined via a TOC analyzer (GB/T 30740–2014). The TN content of the soils was determined using an automatic Kjeldahl apparatus (Liu et al., 2023). Additionally, the determination of soil TP concentration was performed using the Mo-Sb colorimetric method. The soil NH_4_^+^-N and NO_3_L-N contents were measured by indophenol blue colorimetry and vanadium chloride spectrophotometry with a potassium chloride (KCl) extraction method, respectively (Zhang et al., 2019; Zhang et al., 2023b). The soil AP content was measured following the methods of Olsen (1954). Detailed physical and chemical properties of the soils are listed in Table S2.

### 4.3. Determination of 16S rRNA gene amplicon fragments overall and sequence-specific degradation rate constants

The overall and sequence-specific degradation rate constants of added exogenous extracellular 16S rRNA gene amplicon fragments were determined using a primer labeling method, combined with real-time PCR and amplicon sequencing (Fig. 6). Briefly, exogenous eDNA was prepared by PCR amplification using a modified forward primer consisting of a GAPDH F tag fused to the 16S rRNA gene primer 515F, together with the reverse primer 806R. The full primer sequences were as follows: GAPDH-F-515F: 5’-CAT TGG CAA TGA GCG GTT C-GTG CCA GCM GCC GCG GTA A-3’, in which CAT TGG CAA TGA GCG GTT C represents the GAPDH F tag and GTG CCA GCM GCC GCG GTA A represents the 16S rRNA gene forward primer sequence (515F); and 806R: 5’-GGA CTA CHV GGG TWT CTA AT-3’ (Caporaso et al., 2011; Walters et al., 2016). GAPDH is a primer for a human housekeeping gene and it has no homologous sequences in soils. Subsequently, GAPDH was selected as the label primer based on two criteria. First, this primer was selected to avoid interference from the original soil sequences (Huang et al., 2014; Yang et al., 2021; Arvizu-Hernandez et al., 2025), and no detectable PCR amplification was observed for the primer set GAPDH F-806R across all the soil DNA samples included in this study. Second, the melting temperature (Tm) value of GAPDH F approximately matched that of 806R. The GAPDH was incorporated only into the forward primer for several reasons. Methodologically, adding a long linker to the degenerate reverse primer (806R) could reduce amplification efficiency or introduce bias. Economically, single-end labeling allowed us to use the standard reverse primer already carrying sample-specific barcodes, avoiding the costly synthesis of a full set of dual-labeled barcoded primers. This design minimized the risk of secondary structure and primer-dimer artifacts while maintaining sufficient specificity and compatibility with downstream qPCR and sequencing. Subsequently, the exogenous eDNA was separately added to the corresponding fresh soils collected from the 30 study sites, and incubated for 0, 3, 6, 12, 24, and 48 days. Finally, the overall and sequence-specific degradation rate constants of 16S rRNA gene amplicon fragments were calculated based on amplicon sequencing and real-time PCR. Detailed procedures are described as follows.

**Fig. 6.**
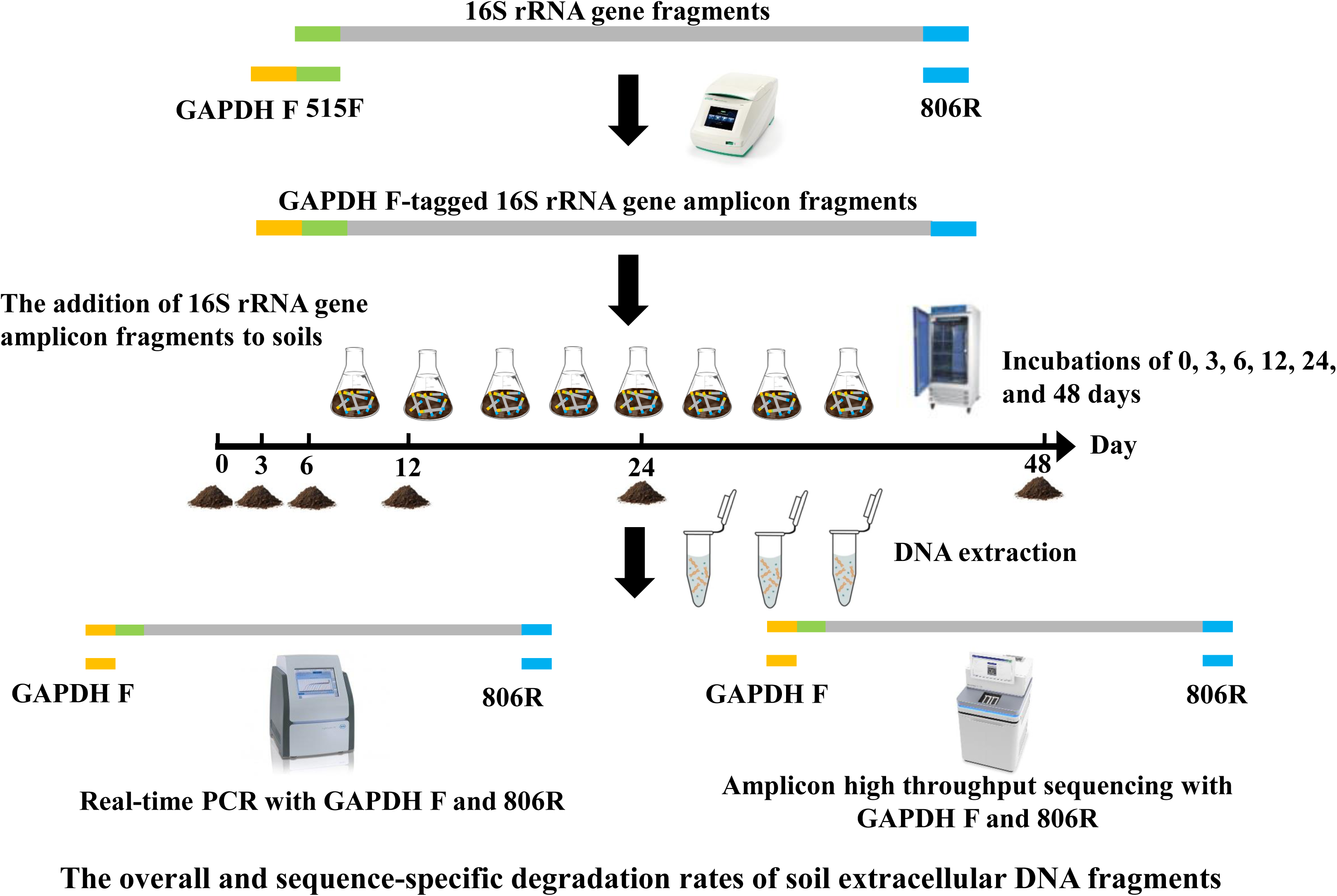
Schematic illustration of the experimental workflow for determining the overall and sequence-specific degradation rates of soil extracellular 16S rRNA gene amplicon fragments.

### 4.4. Preparation of exogenous eDNA

Soil DNA was extracted from 0.5 g of soil using the DNeasy PowerSoil kit (Qiagen, Hilden, Germany) following the manufacturer’s protocols (Li et al., 2023). The GAPDH F-tagged PCR products were generated using DNA extracted from the original soil sample as templates and amplified with GAPDH F-tagged 515F and 806R. The PCR mixture (50 μL) consisted of ExTaq buffer (10×, TaKaRa; 5 μL), dNTP Mix (2.5 mM; 4 μL), forward primer (10 μM; 1 μL), reverse primer (10 μM; 1 μL), template DNA (1 μL), ExTaq (0.25 μL), and DNase-free water (38 μL). The PCR protocol involved an initial denaturation at 95 °C for 10 min, followed by 32 PCR cycles consisting of 30 s at 95 °C, 30 s at 56 °C, and 40 s at 72 °C. Additionally, a final extension was performed at 72 °C for 4 min. Three technical replicates were conducted to amplify each DNA sample, followed by PCR product purification using the GeneJET Gel Extraction Kit (Thermo Scientific, Lithuania). Finally, the purified GAPDH F-tagged 16S rRNA gene fragments were utilized as exogenous extracellular 16S rRNA gene amplicon fragments for subsequent experiments.

### 4.5. Microcosm experiment

The microcosm experiment was conducted using 30 g of soil for each sample. After pre-incubation at 20L for one week, each soil was thoroughly mixed with the GAPDH FLtagged 16S rRNA gene amplicon fragments and incubated further at 20L (Fig. S8). The amount of exogenous GAPDH FLtagged 16S rRNA gene amplicon fragments added to each soil sample was equivalent to 1% of the total DNA concentration naturally present in that soil, as determined fluorometrically prior to the experiment. This concentration was chosen to approximate natural eDNA fluxes resulting from microbial lysis, ensuring experimental relevance to in situ conditions (Table S2). Weekly water additions were performed to maintain the original moisture content of the soils. Soil samples (5 g) were collected after incubation periods of 0, 3, 6, 12, 24, and 48 days with complete mixing prior to each collection. A total of 180 soil samples (30 study sites × 6 incubation times) were included in this study.

We further explored the influence of soil moisture content on the degradation rates of microbial extracellular 16S rRNA gene amplicon fragments based on soils collected from Kaiyuan (KY) and Dashanbao (DSB) in the southwest China. The moisture content gradient included 100%, 75%, 50%, 25%, and 10% of soil water holding capacity. GAPDH FLtagged 16S rRNA gene amplicon fragments originating from the corresponding soils were added to the soils based on the natural soil DNA concentration. The soils were collected after 0, 1, 3, 6, 12, and 24 days of incubation. This complementary experiment included two sites, five moisture levels, six incubation time points, and two replicates per treatment combination, resulting in a total of 120 soil samples.

### 4.6. Soil DNA extraction, real-time PCR, and amplicon sequencing

DNA was extracted from the microcosm experiment soils using the DNeasy PowerSoil kit (Qiagen, Hilden, Germany). The copy numbers of the GAPDH FLtagged 16S rRNA gene amplicon fragments persisting in the soils were determined by employing a LightCycler real-time PCR System (Roche, Mannheim, Germany) with primers GAPDH F and 806R. For each real-time PCR reaction mixture, 1.0 μL of template DNA, 10.0 μL of TB Green™ Premix Ex Taq™ II (Takara, Japan), 0.5 μL of forward primer (20 μM), and 0.5 μL of reverse primer (20 μM) were mixed with 8.0 μL of DNase-free water. The real-time PCR protocol commenced with an initial denaturation at 95 °C for 40 s, followed by 40 cycles (5 s at 95 °C, 30 s at 56 °C, and 40 s at 72 °C). The standard curves exhibited fitted curve *R^2^*values higher than 0.99, with amplification efficiencies of 85%. Each DNA sample quantification was performed in triplicate.

The community profiles of the GAPDH F-tagged 16S rRNA gene amplicon fragments were determined using high-throughput amplicon sequencing. Briefly, GAPDH F-tagged 16S rRNA gene amplicon fragments from the microcosm soils were first amplified from individual samples using GAPDH F and barcode-labeled 806R primers. The reverse primer 806R carried a 12-bp sample-specific barcode, whereas the GAPDH F primer did not contain a barcode. Therefore, each sample was assigned a unique barcode during PCR, which allowed sample demultiplexing after sequencing. The PCR reaction system and thermal cycling conditions were similar to those described above, except that the number of amplification cycles was increased to 35 to obtain sufficient amplicon products for sequencing. The barcoded PCR products from individual samples were purified using a GeneJET Gel Extraction Kit (Thermo Scientific, Lithuania), quantified, and then pooled in equimolar amounts for subsequent library construction. Sequencing libraries were prepared from the pooled barcoded amplicons using the ALFA-SEQ DNA Library Prep Kit according to the manufacturer’s protocol. Universal Illumina-compatible adapters were first ligated to the pooled amplicon products, followed by bead-based purification. An indexing PCR was then performed using the index primer mix, which introduced the complete P5/P7 flow-cell binding sequences and a library-level Illumina index into the pooled library molecules. The indexed library was purified, quantified, and subjected to paired-end sequencing on the NovaSeq platform at MAGIGENE Co., Ltd. (Guangzhou, China).

The USEARCH (v11) with de-noising algorithm was used to analyze the raw sequences (Edgar, 2013). Briefly, paired-end reads were merged using USEARCH, and primer sequences (GAPDH-F-515F and 806R) were removed using the search_pcr2 script. Reads with more than two primer mismatches were discarded. Quality filtering was performed using the fastq_filter script, and sequences with quality scores below 20 were removed. Redundant sequences were dereplicated using the fastx_uniques script. ASVs were generated using the UNOISE3 nonLclustering denoising algorithm (Edgar, 2016), which infers 100% exact sequence variants by distinguishing biological sequences from PCR/sequencing errors. ASVs with total sequence counts fewer than 9 across all samples were removed to reduce noise. To quantify the abundance of each ASV, an ASV table was generated by mapping the qualityLfiltered raw reads back to the ASV set using the otutab command. A 97% similarity threshold was applied for this recruitment to accommodate stochastic sequencing noise while maintaining biological resolution. Crucially, the mapping followed a best-hit priority rule, where each read was assigned to the ASV with the highest percent identity within the 97% radius. This approach ensures that reads derived from the same biological template are accurately counted toward their respective ASV, preventing the underestimation of abundances that would occur with exact matching while strictly preserving the single-nucleotide resolution of the ASV framework. Taxonomic annotation of the ASVs was performed in QIIME2 with the Silva v138 database. A total of 89322 prokaryotic ASVs were obtained. To standardize sequencing depth across samples, the read number of each sample was rarefied to 53251 using the rarefy function in the vegan package in R. The soil prokaryotic diversity (*i.e.,* richness) was assessed via the vegan package in R (Oksanen et al., 2021). All the raw sequencing data and analysis codes have been deposited in the NCBI Sequence Read Archive under BioProject PRJNA1141901 and GitHub repository (https://github.com/lt916/16S-rRNA-genes-rates.git), respectively. The BioSample accession numbers for the 16S rRNA gene amplicon sequencing data are SAMN42909669—SAMN42909908.

### 4.7. Determination of soil extracellular 16S rRNA gene amplicon fragments degradation rate constants

The overall degradation rate constants of the extracellular 16S rRNA gene amplicon fragments were determined by examining the relationships between the copies of labeled 16S rRNA gene amplicon fragments and incubation time, and they were mainly indicated by degradation rate constants. The relationships were fitted using first-order enzyme-catalyzed reaction kinetics, as described by the following equations.

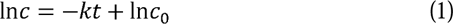

*t*: incubation time, day;

*c*: the copies of GAPDH FLtagged 16S rRNA gene amplicon fragments at time *t*; *c_0_*: the initial copies of GAPDH FLtagged 16S rRNA gene amplicon fragments;

*k*: the degradation rate constants of soil extracellular 16S rRNA gene amplicon fragments, day^-1^;

The sequence-specific degradation rate constants of the exogenous extracellular 16S rRNA gene amplicon fragments were determined using an approach similar to that used for the overall degradation rates of the extracellular 16S rRNA gene amplicon fragments. The absolute abundance of each prokaryotic taxa was estimated by multiplying its relative abundance with the copies of total GAPDH FLtagged 16S rRNA gene amplicon fragments. Subsequently, the reaction rate constants (*k*) were calculated by fitting the relationships between the sequence-specific 16S rRNA gene amplicon fragment copies and incubation time. When comparing the rates of first-order reactions, the reaction rate constant (*k*) usually serves as the core indicator. Therefore, we used the degradation rate constant (*k*) to characterize the degradation dynamics of the added exogenous DNA, which serves as a proxy for how eDNA may be degraded in soils. Because these estimates combine relative abundance profiles with total qPCR abundance, the resulting sequence-specific degradation rates should be interpreted as apparent ASV-level degradation patterns within this analytical framework.

### 4.8. Impacts of eDNA on prokaryotic abundance and diversity analysis

To further explore the implications of the overall and sequence-specific degradation rates of soil extracellular 16S rRNA gene amplicon fragments, we also determined the impacts of extracellular 16S rRNA gene amplicon fragments on soil prokaryotic community analysis and their links with the overall and sequence-specific degradation rates. To inhibit amplification of eDNA, soils were incubated with PMA, as described previously (Carini et al., 2016). Upon photoactivation, eDNA can form covalent bonds through cross-linking, leading to the inhibition of its PCR amplification. In contrast, microbes with intact cell membranes exclude PMA, and their DNA is not cross-linked with PMA, and remains amenable to PCR amplification (Cangelosi and Meschke, 2014). Because cells with intact membranes are less permeable to PMA than eDNA or DNA from membrane-compromised cells, the PMA-treated fraction is expected to be enriched in DNA derived from intact cells. Currently, PMA treatment is widely used to suppress PCR amplification of eDNA (Xue et al., 2023; Canini et al., 2024).

In this study, 0.50 g of soil was mixed with PMA in a total volume of 0.5 mL (40 µM PMA in phosphateLbuffered saline, PBS), while the control soil samples were mixed with PBS without PMA. Both the PMA-treated and control soil samples (PMA-untreated) were gently vortexed for 10 min in the dark at room temperature. Following the incubation, both sets of samples were exposed to a 650 W halogen lamp placed at a distance of 20 cm from the tube, undergoing four consecutive cycles of alternating light (30 seconds) and darkness (30 seconds). Subsequently, the tubes were centrifuged at 10000 × g for 2 min, and the pellets were retained. Finally, DNA was extracted from these pellets using the DNeasy PowerSoil kit (Qiagen, Hilden, Germany) following manufacturer’s protocols.

The abundance of 16S rRNA gene amplicon fragments was determined using quantitative PCR employing a LightCycler real-time PCR System (Roche, Germany). The universal primers set for the 16S rRNA gene amplification were 515F and 806R. The real-time PCR mixture and procedure followed the aforementioned protocol. The 20.0 μL real-time PCR reaction mixture included 1.0 μL of template DNA, 10.0 μL of TB Green™ Premix Ex Taq™ II (Takara, Japan), 0.5 μL of forward primer (20 μM), and 0.5 μL of reverse primer (20 μM) were mixed with 8.0 μL of DNase-free water. The real-time PCR protocol commenced with an initial denaturation at 95 °C for 40 s, followed by 40 cycles (5 s at 95 °C, 30 s at 56 °C, and 40 s at 72 °C). Additionally, prokaryotic 16S rRNA gene fragments were amplified via PCR with universal primers 515F-806R, following the same PCR system, procedure, and bioinformatics analysis as described above. Ultimately, discrepancies in soil microbial properties between the control and PMA-treated samples were utilized to indicate the impact of extracellular 16S rRNA gene amplicon fragments on soil prokaryotic community analysis.

### 4.9. Statistical analysis

The changes in GAPDH F-tagged 16S rRNA gene copies and richness across different incubation time points were determined using repeated-measures analysis of variance. OneLway analysis of variance (ANOVA) followed by Tukey’s honestly significant difference (HSD) postLhoc test was used to compare degradation rate constants among ecosystem types. Prokaryotic community structure differences among the study sites and incubation time points were examined through non-metric multidimensional scaling analysis (NMDS), permutation multivariate analysis of variance (PERMANOVA), and permutational analysis of multivariate dispersion (PERMDISP) (Kruskal, 1964; Anderson, 2001). Random forest modeling was conducted to assess the importance of environmental variables in predicting the overall degradation rates of extracellular 16S rRNA gene amplicon fragments. Structural equation modeling (SEM) was employed to further evaluate the direct and indirect effects of soil moisture, soil pH, MAP, and prokaryotic abundance on the overall degradation rates of extracellular 16S rRNA gene amplicon fragments (Grace, 2006). The sequence-specific degradation rate constants of the exogenous extracellular 16S rRNA gene amplicon fragments were visualized using a heatmap with corrplot packages. To reduce noise arising from unstable detection and unreliable model fitting, only ASVs with relative abundances > 0.01%, present in more than 90% of the study sites, and exhibiting a fitted degradation curve with R² > 0.5 were retained for pairwise sequence-specific degradation rate comparisons. As for the analysis, we performed paired *t*Ltests across all the study sites. Thus, the degradation rates were essentially compared within each site, with both values originating from a same soil sample under identical incubation conditions. A positive *t* value indicates that the first ASV has a significantly higher degradation rate than the second one, and a negative *t* value indicates the opposite. The *p* values were adjusted for multiple comparisons using the FDR method. The relationships between the sequence-specific degradation rate profiles of soil extracellular 16S rRNA gene amplicon fragments and environmental factors were analyzed by Mantel tests (Mantel, 1967).

The impacts of extracellular 16S rRNA gene on the abundance and richness of soil prokaryotic community were determined using the paired *t* test. Differences in the relative abundance of microbial taxa between the total and PMA-treated prokaryotic communities were assessed using Wilcoxon test and the taxa with significant differences were visualized with Graphlan (v1.1.3). Mantel test was employed to further investigate the relationships of PMA-treated or total prokaryotic communities and environmental factors. Pearson correlation test was employed to examine the relationships between the pool sizes and degradation rates of sequence-specific soil extracellular 16S rRNA gene amplicon fragments. Only the abundant taxa, with the proportions exceeding 0.01%, were included in the analysis. For each prokaryotic taxon, its extracellular 16S rRNA gene pool size was calculated as the total 16S rRNA gene copies × the relative abundance of the taxon × (1 – intracellular 16S rRNA gene copies of the taxon/total 16S rRNA gene copies of the taxon). Most of the aforementioned statistical analyses were conducted in R software with a range of packages including vegan, ggplot2, RandomForest, piecewiseSEM, corrplot, matlab, igraph, ggcor, ggpubr, Rmisc, and Hmisc (Breiman, 2001; Cutler et al., 2012; Lefcheck, 2016; Harrell and Dupont, 2020; Oksanen et al., 2021).

## Data availability

All the raw sequencing data have been deposited in the NCBI Sequence Read Archive under BioProject PRJNA1141901. The BioSample accession numbers for the 16S rRNA gene amplicon sequencing data are SAMN42909669—SAMN42909908.

## Declaration of competing interest

The authors declare that they have no known competing financial interests or personal relationships that could have appeared to influence the work reported in this paper.

## Supporting information

Highlights

## Acknowledgements

This work was supported by the National Natural Science Foundation of China (42007035, 42261012, and 32560035), Yunnan Fundamental Research Projects (202301AW070004, 202301BF070001-006, and 202301AT070211), Xingdian Youth Talent Support Program of Yunnan Province (YNQR-QNRC-2018-024, and YNQR-QNRC-2020-087), and Science and Technology Project of Yunnan University for Serving Local Development (YDFWDF202509).

## Authors’ Contributions

R.C. designed research. T.L. performed research. T.L., Z.W., S.Z. and Z.Z. analyzed data. R.C., D.L., X.C. and F.W. revised the paper. T.L. and R.C. wrote the paper. All authors contributed to the preparation of the manuscript. All the authors read and approved the final manuscript.

## Notes

Conflict of Interests: The authors have no conflicts of interest to report.

https://dataview.ncbi.nlm.nih.gov/object/PRJNA1141901

### Competing Interest Statement

The authors have declared no competing interest.

### Summary of Updates

First, the objectives and hypotheses of the study were restated in more detail and clarity. We are restructuring the Introduction to ensure our hypotheses are directly supported by the mechanistic rationale discussed. At the same time, we provided a clearer explanation of the mechanism principles regarding the sequence-specific degradation and the application of the PMA treatment method. To maintain technical accuracy, we have standardized the terminology across the text, specifically replacing the term living community with PMA-treated prokaryotic community while explicitly clarifying the distinctions between extracellular DNA and 16S rRNA gene amplicons. Regarding the methodological rigor, we have integrated a comprehensive description of the microcosm experiments and provided the design rationale for the GAPDH F-tagged primers and fusion primers used in Illumina library preparation. Furthermore, we have addressed the concerns regarding our bioinformatic pipeline, particularly the 97% similarity threshold used for read recruitment. To ensure clarity in data visualization, we have reorganized the figures and updated the legends to explicitly define internal comparative labels, while removing any ambiguous interpretations. Finally, we have added a dedicated section in the Discussion to transparently address technical limitations, including potential PCR and extraction biases, the constraints of using 16S rRNA amplicons as proxies for natural extracellular DNA, and the inherent efficiency limits of PMA treatment in complex soil matrices.

