## Supplementary Materials for "The overall and sequence-specific degradation of soil extracellular DNA fragments: rates and influential factors"

**Table S1.** The geographic and climate information of the sampling sites included in this study.

| Sites | Ecosystems | Latitude (°N) | Longitude (°E) | MAT (℃) | MAP (mm) | Altitude (m) |
| --- | --- | --- | --- | --- | --- | --- |
| S1 | Grassland | 44.186 | 116.465 | 1.86 | 279 | 1083 |
| S2 | Grassland | 41.246 | 111.206 | 2.89 | 300 | 1715 |
| S3 | Grassland | 50.168 | 119.385 | -2.08 | 357 | 524 |
| S4 | Grassland | 37.614 | 101.310 | -0.7 | 469 | 3195 |
| S5 | Grassland | 31.248 | 92.079 | -1.26 | 453 | 4596 |
| S6 | Grassland | 31.274 | 92.111 | -0.24 | 445 | 4486 |
| S7 | Grassland | 30.831 | 91.146 | -1.5 | 418 | 4793 |
| S8 | Grassland | 31.440 | 90.030 | -0.66 | 423 | 4732 |
| S9 | Grassland | 34.838 | 98.324 | -3.35 | 340 | 4215 |
| S10 | Grassland | 36.113 | 100.451 | 3.17 | 364 | 2954 |
| S11 | Desert | 44.291 | 87.923 | 7.46 | 173 | 475 |
| S12 | Desert | 37.016 | 80.724 | 12.83 | 39 | 1362 |
| S13 | Desert | 39.494 | 110.191 | 6.55 | 371 | 1296 |
| S14 | Desert | 37.014 | 80.728 | 12.8 | 40 | 1364 |
| S15 | Cropland | 43.533 | 124.817 | 4.97 | 596 | 203 |
| S16 | Cropland | 36.831 | 116.562 | 13.14 | 605 | 21 |
| S17 | Cropland | 26.762 | 111.863 | 18.14 | 1373 | 117 |
| S18 | Cropland | 30.378 | 112.622 | 16.64 | 1094 | 27 |
| S19 | Cropland | 31.271 | 105.455 | 16.54 | 1001 | 444 |
| S20 | Cropland | 44.296 | 87.931 | 7.47 | 168 | 473 |
| S21 | Forest | 40.420 | 116.661 | 10.79 | 507 | 164 |
| S22 | Forest | 29.800 | 121.783 | 15.8 | 1288 | 170 |
| S23 | Forest | 33.588 | 108.876 | 11.68 | 734 | 732 |
| S24 | Forest | 21.917 | 101.167 | 21.69 | 1626 | 702 |
| S25 | Forest | 36.256 | 108.073 | 8.85 | 499 | 1348 |
| S26 | Forest | 23.052 | 112.473 | 22.53 | 1809 | 13 |
| S27 | Forest | 26.086 | 119.286 | 20.58 | 1382 | 15 |
| S28 | Forest | 35.586 | 118.257 | 12.77 | 778 | 163 |
| S29 | Forest | 38.454 | 106.267 | 8.95 | 204 | 1112 |
| S30 | Forest | 23.381 | 99.421 | 21.41 | 1429 | 991 |

MAT: mean annual temperature and MAP: mean annual precipitation.

**Table S2.** The soil properties of the sampling sites included and the amount of exogenous DNA added in this study.

| Sites | pH | NO_3_^-^-N (mg kg^-1^) | NH_4_^+^-N (mg kg^-1^) | Moisture  (%) | TN  (g/kg) | TP (g/kg) | TOC  (g/kg) | AP  (mg kg^-1^) | Amount  (ng) |
| --- | --- | --- | --- | --- | --- | --- | --- | --- | --- |
| S1 | 7.26 | 19.00 | 3.17 | 6.49 | 1.72 | 0.39 | 15.76 | 22.50 | 747.5 |
| S2 | 6.69 | 10.47 | 3.73 | 3.67 | 1.45 | 0.36 | 14.92 | 12.71 | 480 |
| S3 | 6.71 | 14.30 | 7.53 | 7.31 | 2.76 | 0.50 | 33.12 | 20.61 | 1442.5 |
| S4 | 6.55 | 55.85 | 3.27 | 35.67 | 4.19 | 0.74 | 44.50 | 26.35 | 2835 |
| S5 | 6.72 | 58.82 | 2.88 | 29.21 | 4.36 | 0.71 | 51.28 | 18.09 | 1682.5 |
| S6 | 6.69 | 53.09 | 2.71 | 27.71 | 4.07 | 0.58 | 47.81 | 14.50 | 1250 |
| S7 | 6.42 | 44.72 | 3.04 | 10.42 | 2.70 | 0.53 | 28.09 | 14.14 | 857.5 |
| S8 | 6.21 | 77.65 | 2.55 | 12.8 | 2.37 | 0.47 | 22.05 | 23.12 | 532.5 |
| S9 | 8.85 | 10.03 | 2.17 | 7.60 | 0.78 | 0.36 | 7.41 | 35.94 | 417.5 |
| S10 | 8.16 | 5.96 | 3.22 | 6.51 | 0.33 | 0.27 | 4.28 | 18.14 | 265 |
| S11 | 7.83 | 30.12 | 3.42 | 2.89 | 0.66 | 0.97 | 7.25 | 62.48 | 245 |
| S12 | 8.31 | 19.56 | 2.76 | 1.29 | 0.14 | 0.50 | 2.27 | 12.76 | 197.5 |
| S13 | 8.34 | 3.58 | 2.96 | 0.76 | 0.54 | 0.45 | 10.38 | 84.64 | 200 |
| S14 | 8.07 | 22.89 | 10.96 | 0.77 | 0.24 | 0.61 | 3.75 | 85.99 | 255 |
| S15 | 5.98 | 17.93 | 2.88 | 14.83 | 1.13 | 0.39 | 18.57 | 57.94 | 535 |
| S16 | 6.14 | 28.99 | 2.55 | 15.16 | 1.17 | 1.00 | 10.79 | 42.66 | 1110 |
| S17 | 6.03 | 52.11 | 4.12 | 23.41 | 1.99 | 0.56 | 21.83 | 28.50 | 1045 |
| S18 | 6.23 | 43.84 | 2.81 | 18.59 | 1.60 | 0.74 | 17.96 | 35.32 | 495 |
| S19 | 6.98 | 21.37 | 3.94 | 15.72 | 0.68 | 0.51 | 8.49 | 24.19 | 314 |
| S20 | 6.73 | 14.18 | 2.73 | 10.60 | 0.71 | 1.18 | 7.19 | 81.96 | 475 |
| S21 | 7.10 | 14.74 | 4.35 | 18.88 | 1.10 | 0.42 | 22.45 | 19.15 | 897.5 |
| S22 | 6.84 | 7.19 | 4.14 | 22.48 | 0.43 | 0.18 | 5.34 | 9.74 | 275 |
| S23 | 7.33 | 137.54 | 3.58 | 35.22 | 4.73 | 0.90 | 58.93 | 26.87 | 2737.5 |
| S24 | 6.12 | 61.38 | 55.55 | 26.29 | 2.33 | 0.37 | 22.89 | 29.58 | 1352.5 |
| S25 | 7.70 | 58.99 | 2.65 | 15.65 | 1.33 | 1.15 | 11.74 | 29.56 | 957.5 |
| S26 | 4.15 | 15.94 | 34.82 | 16.54 | 2.13 | 0.28 | 30.01 | 15.58 | 632.5 |
| S27 | 5.46 | 56.22 | 2.73 | 29.77 | 1.97 | 1.15 | 23.72 | 13.78 | 1197.5 |
| S28 | 5.22 | 63.38 | 18.82 | 16.03 | 0.99 | 0.43 | 8.11 | 149.84 | 292.5 |
| S29 | 4.81 | 147.32 | 2.81 | 48.78 | 3.69 | 0.62 | 54.98 | 43.58 | 905 |
| S30 | 4.98 | 22.94 | 2.88 | 28.82 | 0.99 | 0.32 | 10.50 | 18.45 | 410 |

NO_3_^-^-N: soil NO_3_^-^-N contents; NH_4_^+^-N: soil NH_4_^+^-N contents; TN: soil total N contents; TP: soil total P contents; AP: soil available P contents; and TOC: soil total organic carbon contents.

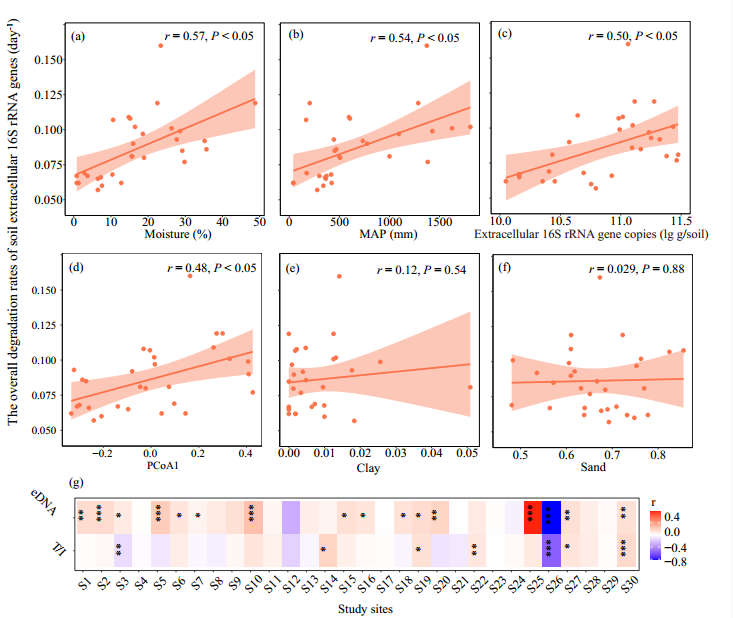

**Fig. S1.** (a)–(f) The relationships between the overall degradation rates of soil extracellular 16S rRNA gene amplicon fragments and environmental factors; (g) the relationships among the sequence-specific degradation rates, contents, and the influencing intensity of extracellular 16S rRNA gene amplicon fragments. The eDNA means the copies of extracellular 16S rRNA gene amplicon fragments. The T/I represents the ratios between the relative abundance of each prokaryotic ASVs based on the total DNA extraction to that based on intracellular DNA extraction, and it is used to indicate the influencing intensity of extracellular 16S rRNA gene amplicon fragments on the relative abundance of prokaryotic taxa. Moisture: soil moisture content; MAP: mean annual precipitation; and NMDS1: the scores at the first axis of the NMDS ordination of prokaryotic community profile.

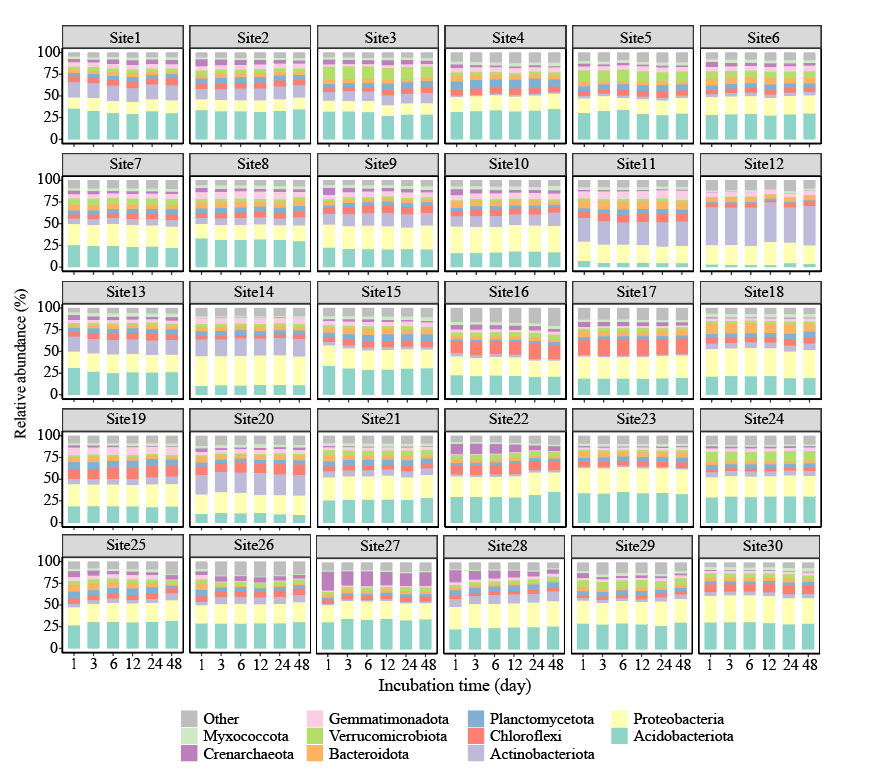

**Fig. S2.** Soil prokaryotic community composition based on GAPDH F‑taggedGAPDH F‑tagged 16S rRNA gene amplicon fragments at different incubation time points.

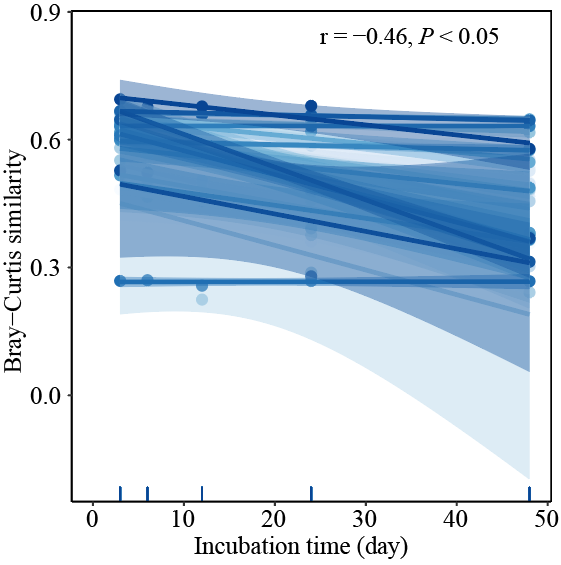

**Fig. S3**. The relationships between community profiles similarities based on the GAPDH F‑tagged 16S rRNA gene amplicon fragments and the intervals of incubation time. The similarities were calculated between the community profiles at time 0 and the other time points.

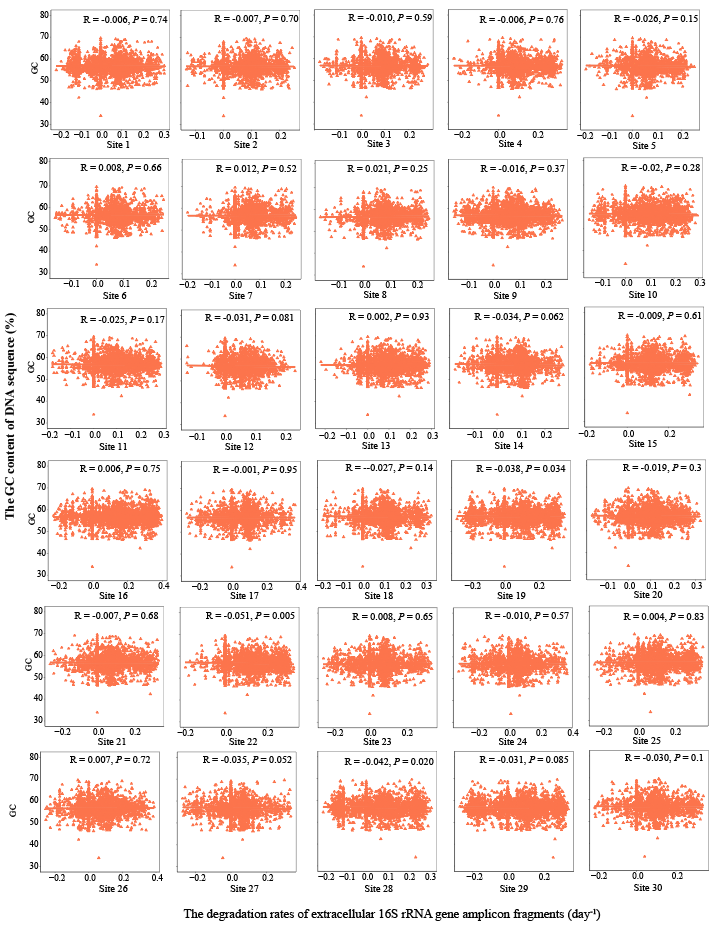

**Fig. S4**. The relationships between the degradation rates of extracellular 16S rRNA gene amplicon fragments and the G+C content of DNA sequence.

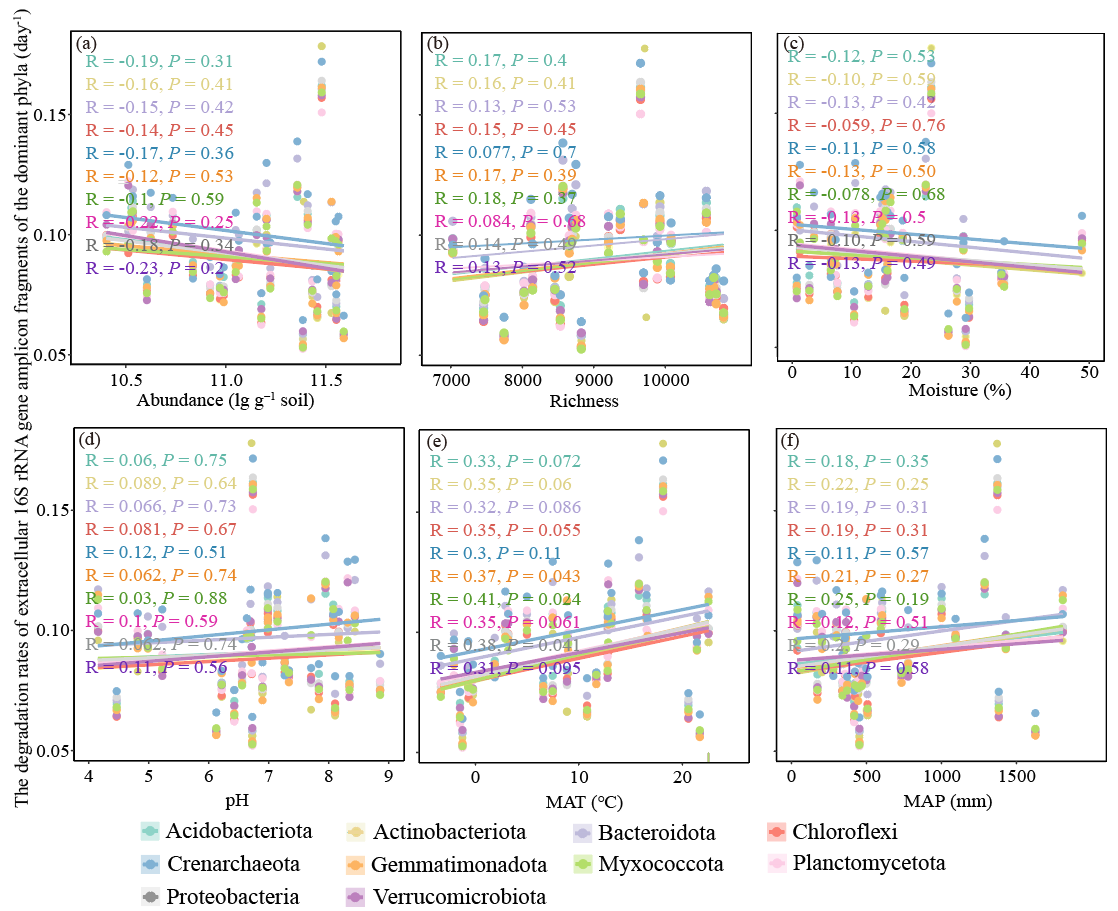

**Fig. S5**. The relationships between the degradation rates of extracellular 16S rRNA gene amplicon fragments of the dominant phyla (top 10) and different influencing factors. MAT: mean annual temperature and MAP: mean annual precipitation.

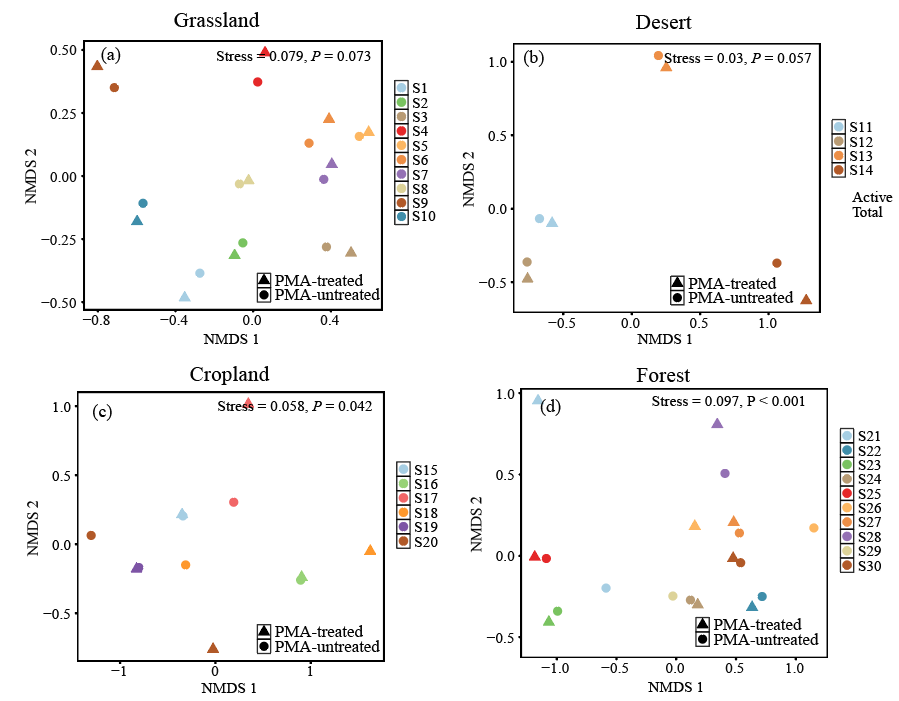

**Fig. S6.** The nonmetric multidimensional scaling (NMDS) ordination of total and PMA-treated soil prokaryotes in different ecosystems. The different colors represent samples from different sites;

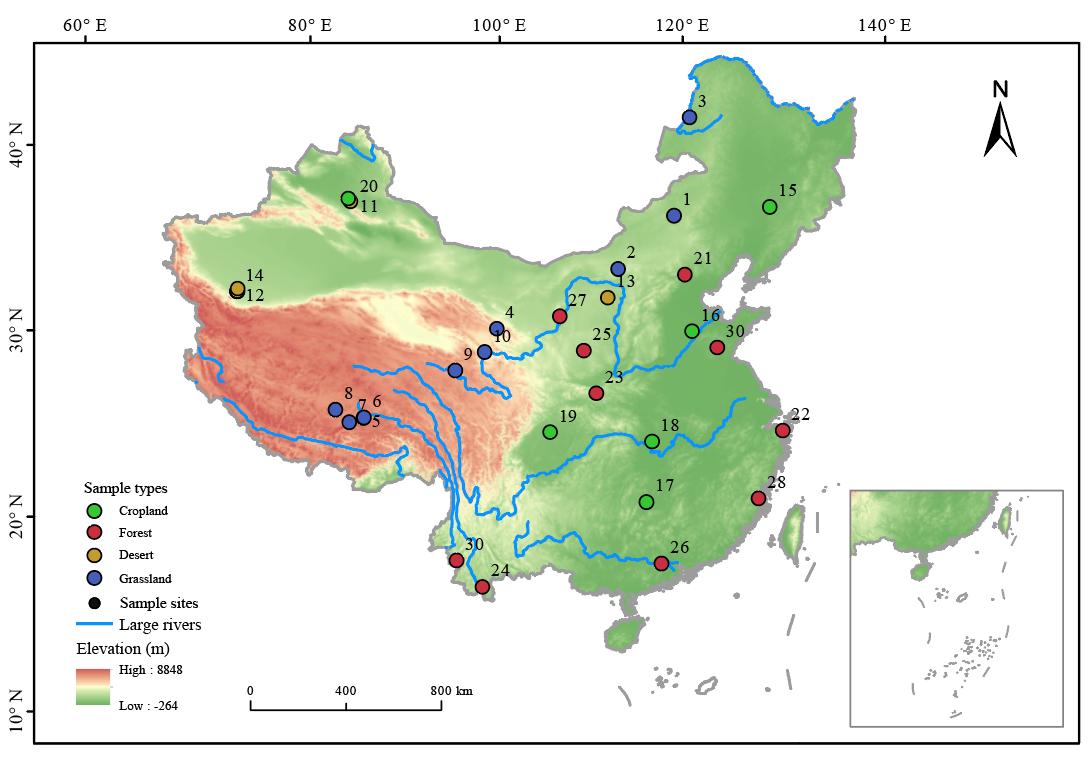

**Fig. S7**. The map of the sampling sites.

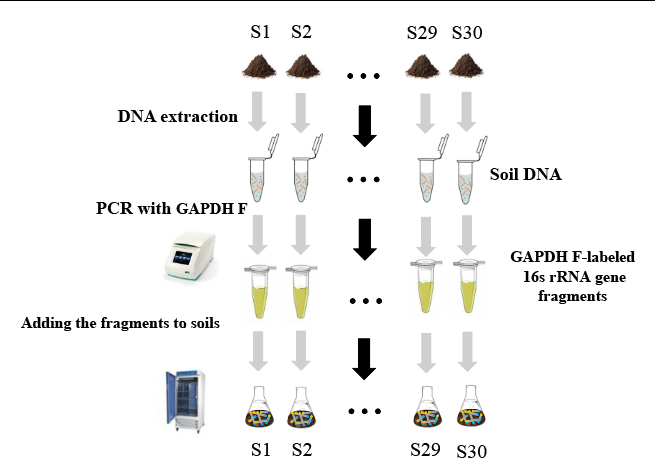

**Fig. S8**. The experimental design.
