## Supplementary material for "The overall and sequence-specific degradation of soil extracellular DNA fragments: rates and influential factors": Highlights

Overall and sequence-specific degradation rate constants of soil eDNA fragments were determined.

Soil extracellular 16S rRNA gene overall degradation rates highly varied across China.

The sequence-specific degradation rates of eDNA fragments substantially varied.

The sequence -specific degradation of eDNA largely affected soil microbial analysis.

Soil eDNA degradation was mainly influenced by water availability.
